# EGCG modulates nuclear formaldehyde-induced Tau phosphorylation in Neuronal cells

**DOI:** 10.1101/2020.10.04.325134

**Authors:** Shweta Kishor Sonawane, Anshu Raina, Amitabha Majumdar, Subashchandrabose Chinnathambi

**Affiliations:** Neurobiology Group, Biochemical Sciences Division, CSIR-National Chemical Laboratory, Dr. Homi Bhabha Road, Pune-411008, India; Academy of Scientific and Innovative Research (AcSIR), Pune-411008, India; Drosophila Neurobiology lab, National Centre for Cell Science, Pune-411007, India

**Keywords:** Alzheimer’s disease, Nuclear Tau, Tau phosphorylation, CDK5, Drosophila model of Tau phosphorylation, EGCG

## Abstract

Tau hyperphosphorylation is one of the major causes of Alzheimer’s disease pathology. The abnormal phosphorylation curtails the physiological function of Tau of microtubule stabilization and renders it more prone to aggregation. Apart from its function in the cytoplasm, Tau is attributed to play a role in the nucleus. Nuclear function of Tau is dependent on its residue-specific phosphorylation. We studied the effect of a green tea polyphenol, EGCG, on the formaldehyde-induced Tau phosphorylation and Tau kinase CDK5. Interestingly, we observed unique localization of phospho-Tau (AT 8 and AT 100) in the nucleus in various EGCG treatments. EGCG was also found to lower the levels of CDK5 in the formaldehyde-treated cells. Further, the role of EGCG was tested *in vivo* in drosophila eye model of hyperphosphorylated Tau (Tau E14). The results suggest that EGCG can modulate nuclear Tau phosphorylation and lower the levels of Tau kinase CDK5.

## Introduction

Tau pathology in Alzheimer’s disease is known to hamper several cellular functions including microtubule dynamics, proteostasis *etc*. Tau is a microtubule-stabilizing protein, playing an important role in maintaining the integrity of the axonal microtubules ^1,2^. Tau was thought to be a cytosolic protein but additional roles of Tau in various sub-cellular compartments have also been reported ^3^. Tau is present in the nucleus and known to protect DNA against heat-induced DNA damage. Intense oxidative stress and mild heat stress leads to build-up of dephosphorylated Tau in the nucleus of the neurons ^4^. *In vitro*, Tau interacts preferentially with AT-rich region of DNA ^5^. Though, Tau is known to be localized to nucleus, the exact pathways and signals required for this are not fully understood. Recently various studies have been reported that nuclear Tau is preferentially phosphorylated at specific residues. This sheds some light on Tau phosphorylation and its localization to the nucleus ^6,7^. Tau phosphorylation has been explored extensively; hyperphosphorylation of Tau is known to be involved in aberrant Tau aggregation. Hyperphosphorylated Tau carries 8 molecules of phosphates per molecule whereas; physiologically Tau carries 2 phosphates per molecule ^8^. Phosphorylation of Tau is detected by various epitope specific antibodies such as AT8 (S202/T205), AT100 (S212/T214), AT180 (T231), AT270 (T181), PHF-1 (S396/S404), PHF-6 (T231) ^9,10^. Thus, these epitopes also serve as the diagnostic markers for AD. Most of these epitopes are sites of proline-directed kinases-like GSK3β, CDK5, MAPK *etc*.

CDK5 is a non-classical cyclin-dependent kinase, which plays a major role in Tau phosphorylation. CDK5 is predominantly localized in the nucleus of the post-mitotic neurons and helps to maintain this state ^11^. CDK5 is a proline-directed ser/thr kinase and plays a crucial role in Tau pathology by abnormally hyperphosphorylating Tau ^12^. CDK5 phosphorylates 9-13 sites in Tau ^13^. Ser 202 and Thr 205 are the exclusive sites for CDK5 phosphorylation ^14^. The abnormal phosphorylation of Tau by CDK5 is more pronounced because the sites phosphorylated by CDK5 are more resistant to dephosphorylation as compared to phosphorylation by PKA ^15^. Thus, CDK5 localization and its Tau phosphorylation play a major role in the pathogenesis of AD ^16,17^.

EGCG is reported to reduce the level of phosphorylated Tau in primary cortical neurons and inhibit Tau aggregation ^18^. EGCG has multiple targets in neuroprotection. It decreases the cellular ROS ^19^ and inhibits the expression of cell death and pro-inflammatory genes ^20,21^. Thus, EGCG acts as neuroprotective molecules by modulating multiple pathways. We report the role of EGCG in modulating two phospho-Tau epitopes AT 8 and AT 100. We also used *Drosophila melanogaster* as a model to exhibit the effect of EGCG on neurodegeneration caused by Tau protein. The model employed for this purpose is Elav GMR hTau, which expresses the 2N4R isoform of human Tau protein in Drosophila compound eye thus, providing a means for distinct qualitative evaluation of neurodegeneration ^22^. Though, EGCG did not show any toxic symptoms in fly model, we did not observe distinct variation in Elav GMR human Tau model after treatment.

## Results

### EGCG alters CDK5 levels in neuronal cells

CDK5 is a prime proline-directed serine/threonine kinase, which is functionally active in neurons ^23,24^. CDK5 is implicated in Tau hyperphosphorylation.. We show that EGCG alters the levels CDK5 (Fig. 1A). Amongst the 6 treatment groups including cell control, EGCG, EGCG pretreatment and simultaneous treatment showed reduced levels of CDK5 whereas; Formaldehyde (FA) treated cells showed enhanced levels. Though the other treatment groups did not show a significant reduction in CDK5 levels, the levels were maintained same as control (Fig. 1B, S1).

**Figure 1.**
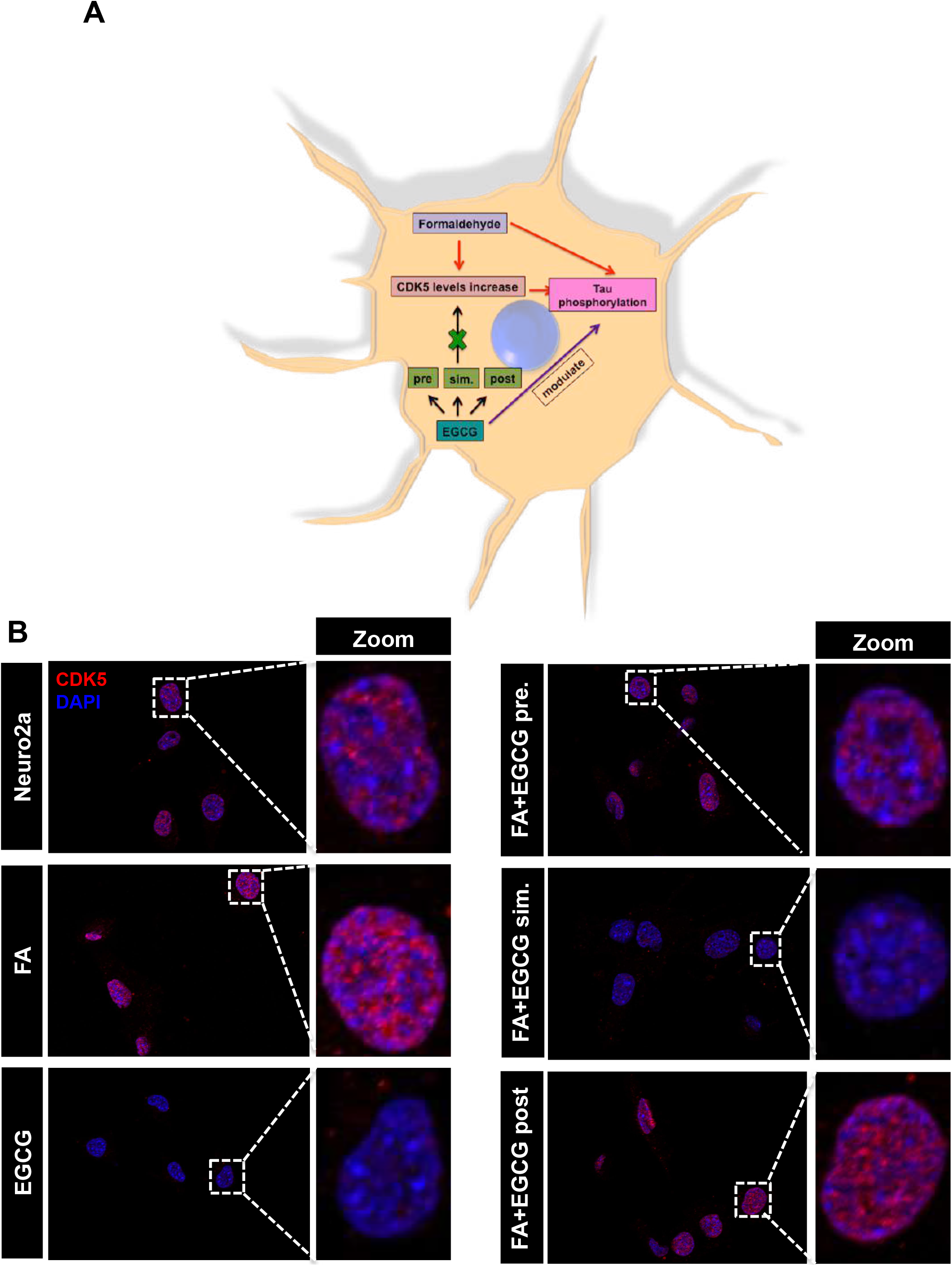
EGCG alters CDK5 levels in neuronal cells. A) Hypothesis suggesting increase in CDK5 levels on formaldehyde treatment followed by Tau phosphorylation. EGCG treatment might reduce this Tau kinase level in turn modulating Tau phosphorylation. B) Control shows basal level of CDK5 in neuro2a cells whereas FA treated cells show enhanced CDK5 level as compared to control. Treatment of EGCG alone and previous to FA and simultaneously with FA showed reduced levels of CDK5 as compared to positive control. EGCG and post treatment did not show effect in altering the CDK5 as compared to positive control.

### EGCG modulates FA-induced Tau phosphorylation

In order to study effect of EGCG on formaldehyde-induced Tau phosphorylation, we studied two epitopes with dual phosphorylation, AT8 (pS202/pT205) and AT100 (pS212/pT214) for the same treatment groups as for CDK5. The immunofluorescence for AT8 showed enhanced and distinct nuclear localization in the nucleus in the pretreatment group (Fig 2A, B). Simultaneous and post-treatment groups showed reduced AT8 signal as compared to positive control (FA treatment). The levels of total Tau remain unchanged in all the groups. In order to study overall phosphorylation, western blotting was carried out for AT8 and AT100 phospho-Tau epitopes. The AT8 phospho-Tau was detected only in control cells at smaller molecular weights rather than at expected molecular weight of 50 kDa (Fig 2C and S2). The loading control actin did not show much variation between the treatment groups. AT100 showed reduced levels in all treatment groups as compared to positive control. EGCG treated group showed more reduction as compared to positive control. In EGCG pretreated group, AT100 showed enhanced fluorescence in the nuclear and cell periphery whereas; the total Tau fluorescence was observed uniformly in the cell except nucleus (Fig. 3A, B). The immunoblotting for AT100 again yielded the signal at lower molecular weights but prominently in control and EGCG treated cells (Fig 3C). In order to check for the distribution of phospho-Tau in nucleus and cytoplasm orthogonal cross-sections were obtained form 3D Z-stacks. AT8 staining was uniform in the control and cells treated with EGCG. Treatment with FA and FA+EGCG (pre-, post, sim-) showed increased localization to nucleus (Fig. S3). AT100 on the other hand showed increased nuclear localization in control and treated cells except post treatment group, which showed cytoplasmic localization of AT100 Tau (Fig. S4). The morphological analysis of cells showed loss of morphology in FA treated cells as opposed to untreated of EGCG treated groups (Fig. S5). Thus, EGCG alters CDK5 levels and modulates Tau phosphorylation.

**Figure 2.**
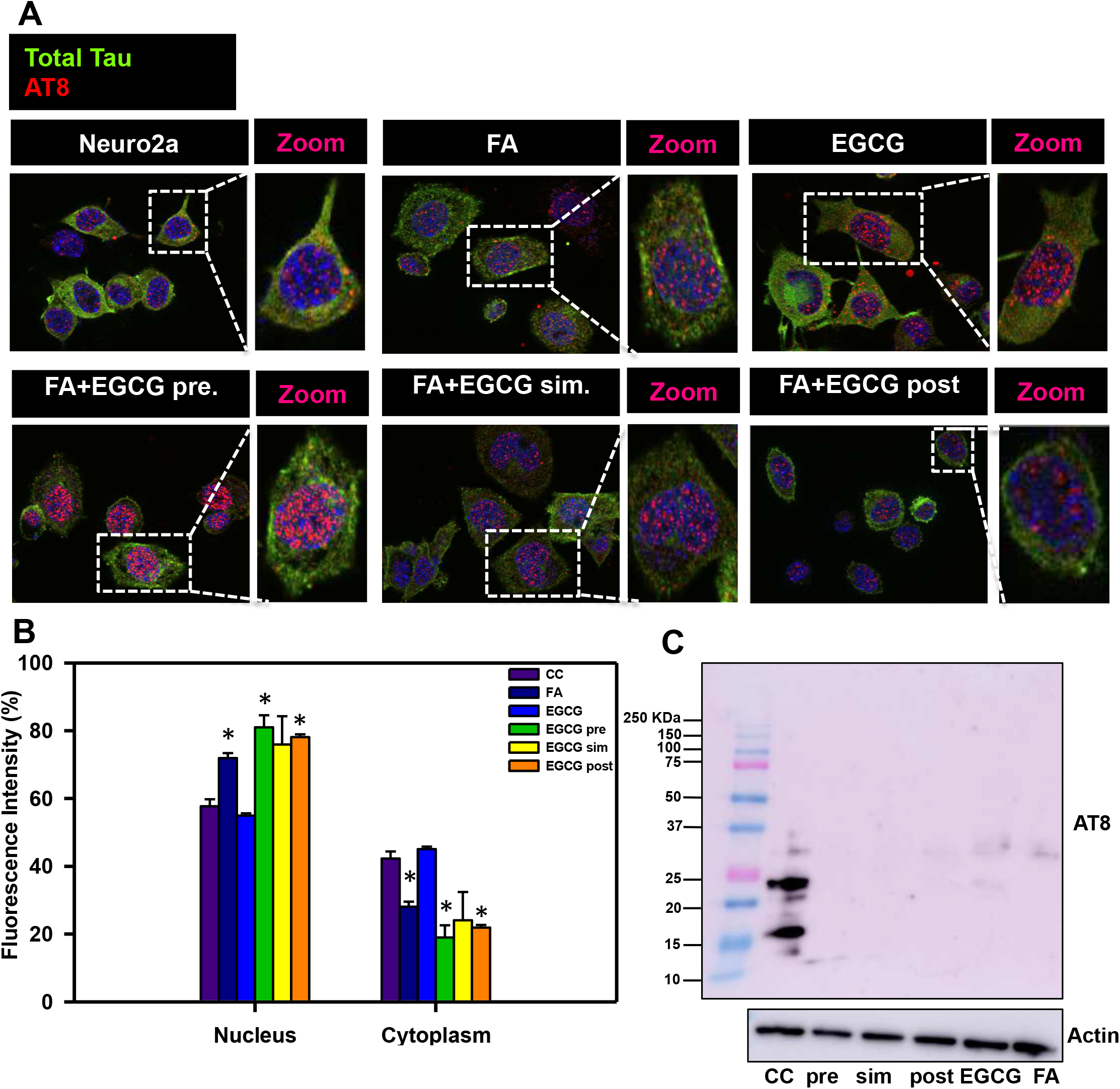
EGCG modulates Tau phosphorylation at AT8 (pS202/pT205) epitope. A) AT8 positive Tau is seen mainly concentrated in the nucleus but is present in cytoplasm as well. The untreated control showed basal level of fluorescence for AT8. EGCG pretreated cells showed enhanced AT8 signal in the nucleus as compared to positive control. Total Tau levels remained same in all the treated cells. B) The quantification reveal that EGCG pretreatment shows more AT8 positive Tau in the nucleus as compared to cytoplasm p<0.05. C) The immunoblotting for AT8 epitope suggests degraded Tau products detected only in control cells.

**Figure 3.**
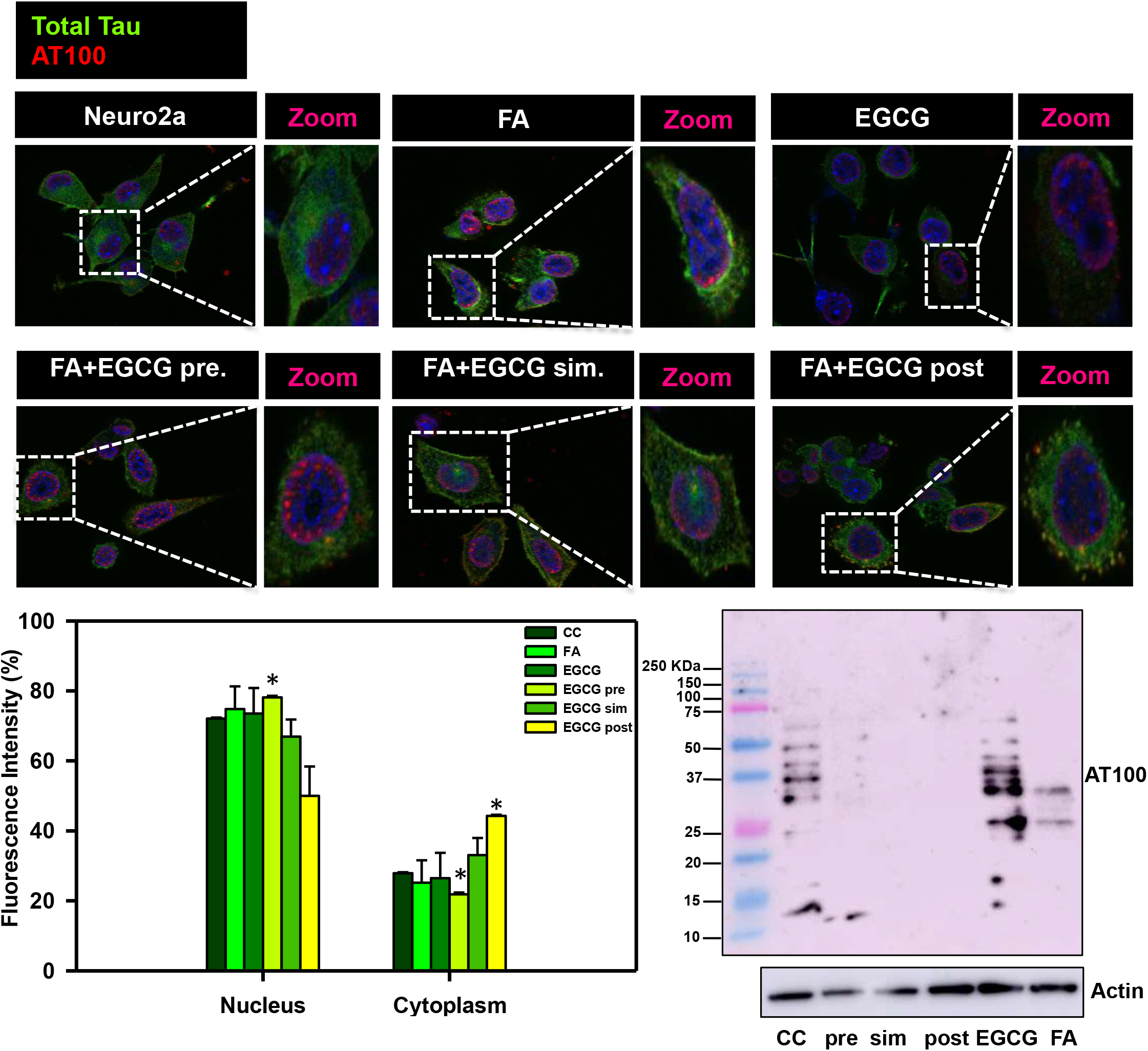
EGCG modulates Tau phosphorylation at AT100 (pS212/pT214) epitope. A) AT 100 positive Tau is observed distinctly in the cell and nuclear periphery. EGCG treated cells showed a reduced AT100 fluorescence as compared to positive control. Other treatment groups did not show change in AT100 fluorescence as compared to positive control. B) The fluorescence quantification reveal that EGCG pretreatment shows more AT100 positive Tau in the nucleus as compared to cytoplasm p<0.05. C) The western blot for AT100 epitope reveals degraded Tau only in control and EGCG treated cells.

### EGCG maintains viability of drosophila model of Tauopathy

In order to check the *in vivo* toxicity of EGCG, the survival assay was carried on drosophila model of Tauopathy. The flies were grown in fly food supplemented with EGCG at 10, 25 µM and water as a vehicle control. The survival assay was carried out for 31 days. 25 flies were analyzed from each group and plotted as percentage alive and dead. It was observed that EGCG treatment showed 96% survival in drosophila and vehicle control showed 92% survival rate (Fig.4). Thus, EGCG was found to be non-toxic *in vivo*.

**Figure 4.**
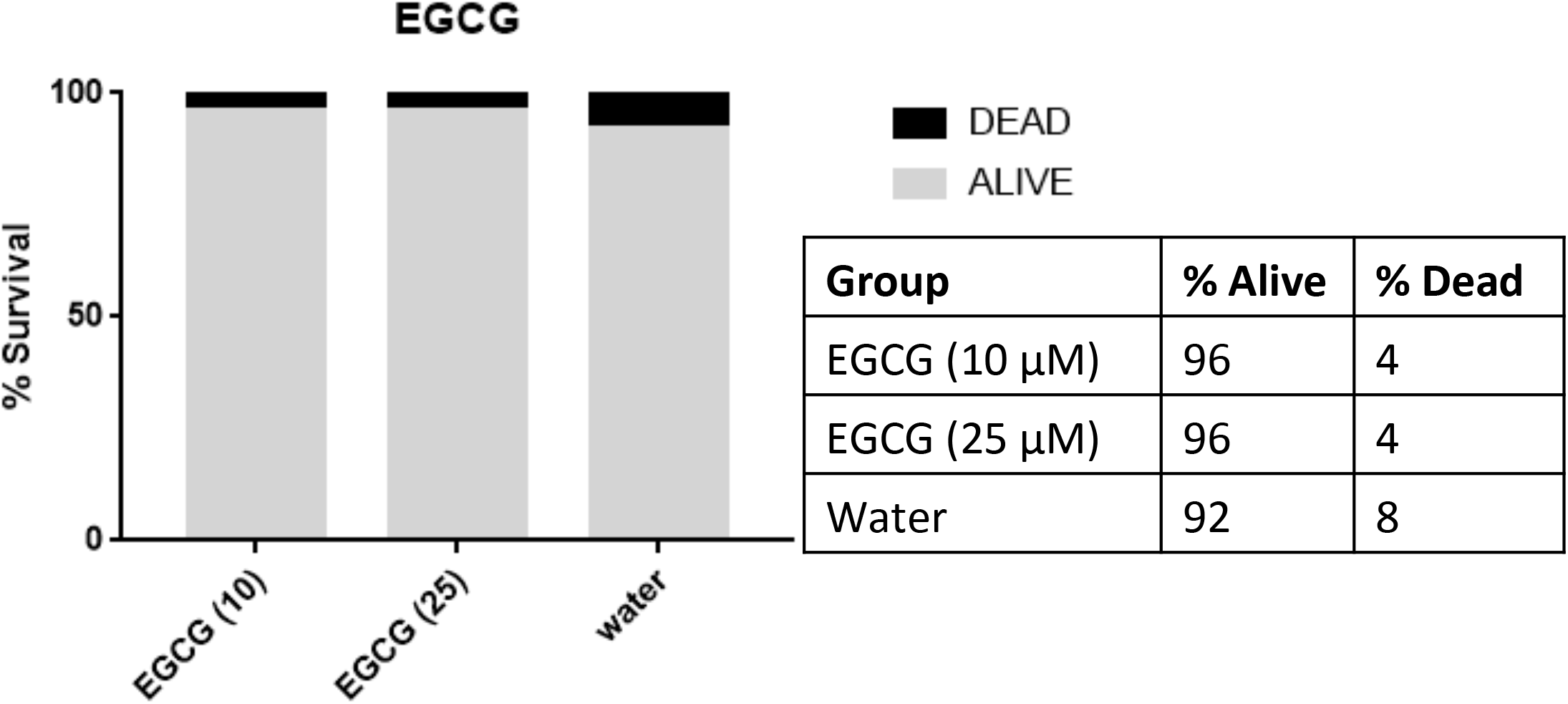
EGCG is not toxic to *Drosophila* model of Tauopathy. The effect of EGCG was studied on fly model of Tauopathy. The survival assay was conducted for 10 and 25 µM of EGCG with water as a vehicle control. EGCG at both the concentrations did not show toxicity in the flies till 31 days. Both the concentrations of EGCG showed 95% survival rate.

### EGCG does not rescue eye phenotype in drosophila model of Tauopathy

EGCG was not found to be toxic to drosophila model, so highest concentration (25µM) was considered for further experiments. We studied the effect of EGCG on the eye phenotype-using drosophila model Tau in the eye a Gal4-UAS system. Expression of Tau leads to rough eye phenotype in drosophila pertaining to marked neurodegeneration. The eye phenotype was monitored for a month and representative images were taken at day 1, 10 and 15 in presence of EGCG and vehicle control water. We did not observe any recue of eye phenotype by EGCG at 25 µM concentration (Fig. 5).

**Figure 5:**
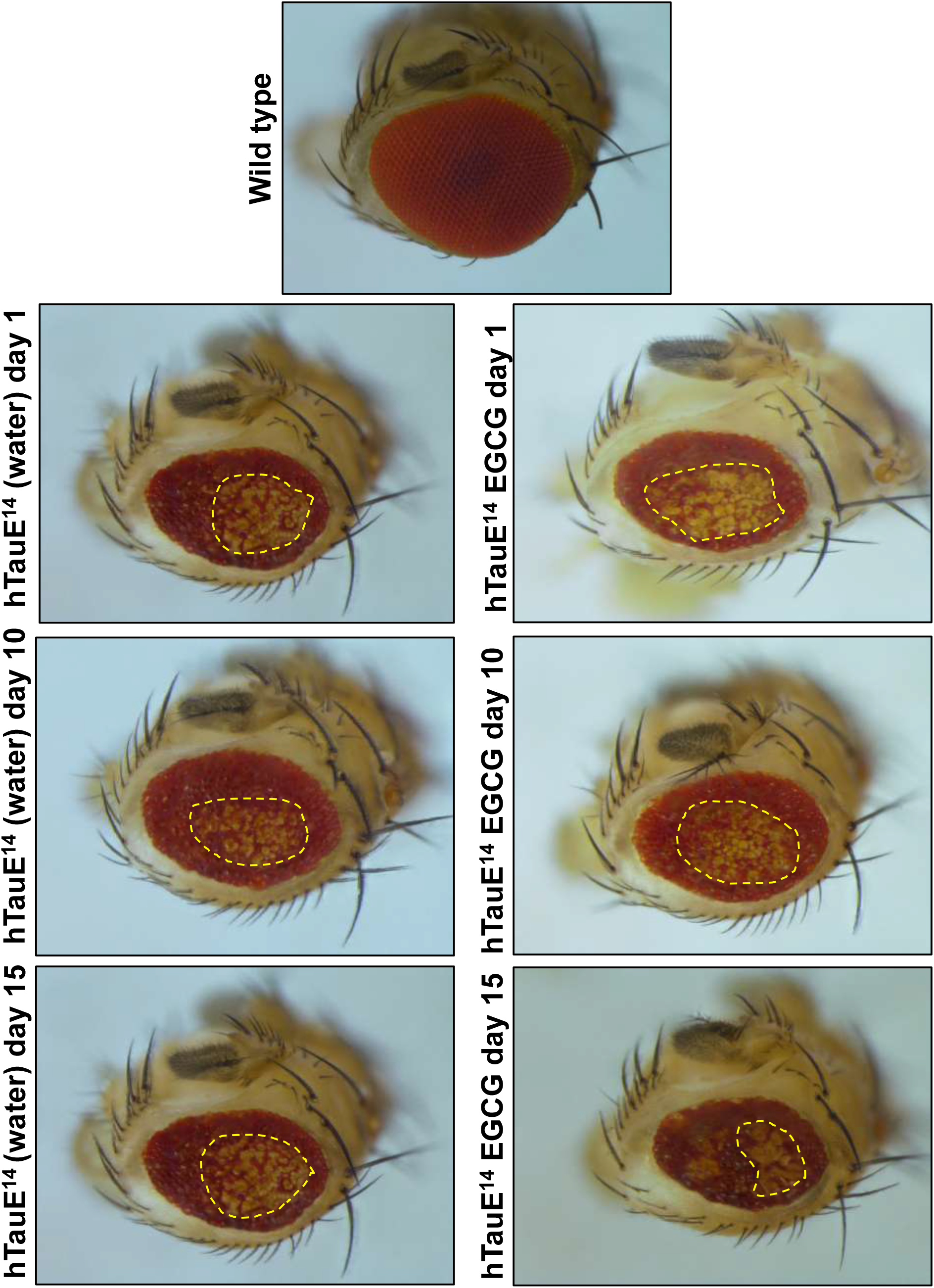
Effect of EGCG on drosophila eye phenotype. The wild type drosophila (Canton S) shows smooth eye phenotype whereas the expressing flies show rough eye phenotype. The control as well as EGCG treatment did not show rescue of eye phenotype in the drosophila model.

## Discussion

Alzheimer’s disease has been attributed to many neuronal pathologies and Tau plays an important role in these pathologies. Tau leads to array of pathologies, which include aggregation and accumulation, cytoskeleton dysfunction, synaptic malfunction etc. One of the important causes of Tau pathology is its excessive abnormal phosphorylation by several kinases. CDK5 is one among the many kinases that hyperphosphorylates Tau and aggravates the aggregation and accumulation. Tau has 16 serine/threonine-proline sites of which 9-13 sites are phosphorylated by CDK5. The major sites of Tau phosphorylated by CDK5 include S202/T205 commonly known as AT8 epitope, S204 and S235 ^25^. Though the other sites include T181, S212, T214 *etc*^26^. EGCG is known to reduce Tau phosphorylation in primary neurons at specific epitopes ^18^Along with Tau hyperphosphorylation, CDK 5 is also implicated in inflammatory responses induced by Aβ oligomers. The administration of roscovitine, a CDK5 inhibitor reduced the inflammatory processes induce by Aβ oligomers, thus, highlighting the role of this kinase in neuroinflammation ^27^. Hence, reduction of CDK5 can not only prevent Tau pathology but also inhibit the inflammatory processes involved in neurodegeneration ^28^. The study of AT 100 nuclear localization has been linked to cell senescence. It has been reported that AT 100 nuclear localization increases with aging. Moreover, the heterochromatin localization of AT 100 has been observed to increase in the early stages but decreases at the later AD stages ^29^. Our study reports decrease in the levels of AT 100 phosphorylated Tau in EGCG treatment groups suggesting delayed aging as compared to formaldehyde treated cells. AT8 Tau on the other hand is associated with neuronal cell differentiation playing a role in regulation of inactive rDNA during differentiation and also a marker o differentiated status of neuronal cells ^30^. Our immunofluorescence studies suggested enhanced nuclear localization of AT8 in EGCG alone and EGCG pretreatment group suggesting its role in inducing neuronal differentiation. EGCG has been reported to enhance the survival of fly model of Parkinson’s disease ^31^. Moreover, EGCG was found to improve fitness and increase fly lifespan by modulating the glucose metabolism ^32^. Thus, EGCG is found to be neuroprotective and enhance survival of various disease models.

## Conclusions

The current study highlights a novel role of EGCG in modulating Tau phosphorylation. EGCG reduced the levels of Tau kinase CDK5. It modulated the localization of nuclear phosphorylated Tau epitopes suggesting potential roles in aging and neuronal differentiation (Fig. 6). Moreover, EGCG could not rescue the eye phenotype in drosophila model of Tauopathy at the given concentration; it did not hamper the fly survival.

**Figure 6.**
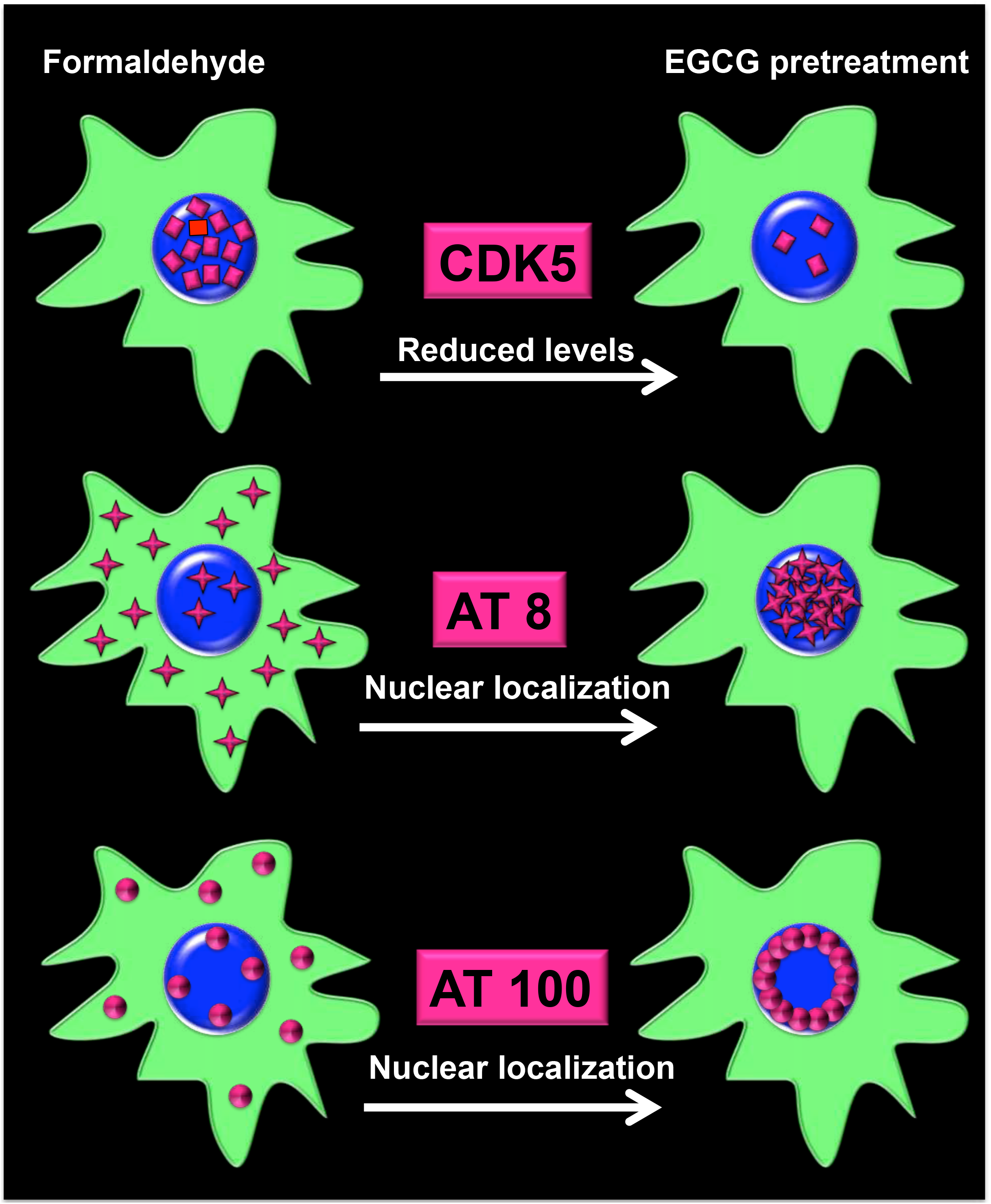
EGCG pretreatment modulates Tau phosphorylation. The cartoon depicts the effect of EGCG pretreatment on the levels of Tau kinase, CDK5 and localization of phosphorylated nuclear Tau. EGCG pretreatment reduces the levels of CDK, which are increased in formaldehyde treatment. Further, it modulates localization of Tau phosphorylated at two epitopes towards nucleus. This nuclear localization may have various implications in AD.

## Materials and methods

### Chemicals and reagents

Formaldehyde, Glycerol and paraformaldehyde were purchased from MP Biomedicals. EGCG was purchased from Sigma-Aldrich. All the cell culture reagents like Advanced DMEM, Fetal Bovine Serum (FBS), Penstrep antibiotic cocktail, Phosphate Buffered Saline (PBS) were purchased from Invitrogen. Antibodies were obtained from respective sources. CDK5 mouse monoclonal (Invitrogen, cat. no. AHZ0492) (1:100), AT8 (Thermo fisher, cat. no. MN1020) AT 100 (Thermo fisher, cat. no. MN1060). Alexa flour-488 (Invitrogen, cat no A-11001). Goat anti-Rabbit IgG (H+L) Cross-Adsorbed Secondary Antibody with Alexa Fluor 555 (A-21428).

### Immunofluorescence

The immunofluorescence studies were carried out by on neuro2a cells. 5×10^4^ cells were seeded on 18 mm glass coverslips. The cells were incubated for 24 hours in Advanced DMEM containing antibiotics and 10% fetal bovine serum. The effect of EGCG on Tau kinase and phosphoepitopes was studied in 6 experimental groups. Formaldehyde (FA) was used as inducer of phosphorylation. The groups comprised of FA only treatment at 0.5 mM; EGCG only group at 10 µM; and EGCG pretreatment + FA group; EGCG + FA group simultaneous treatment; EGCG post-treatment + FA group. The pre- and post-treatment with EGCG was given for 30 minutes. After 4 hours of incubation, cells were washed with cold PBS and fixed with 4% paraformaldehyde. Cell were pemeabilized with 0.2% TritonX 100 and blocked in 2% horse serum for 1 hour. The respective primary antibodies like CDK5 mouse monoclonal (1:100), AT8 (1:100), AT 100 AT100 (1:100) were added onto the cells and incubated in the moist chamber at 4°C overnight. Next day, cells incubated with respective antibodies like anti-mouse secondary antibody conjugated with Alexa fluor 488 (1:1000), Goat anti-Rabbit IgG (H+L) Cross-Adsorbed Secondary Antibody with Alexa Fluor 555 (1:500). The coverslips were further counterstained with DAPI and mounted in 80% glycerol. The imaging was done in in Axio Observer 7.0 Apotome 2.0 (Zeiss) under 63X oil immersion lens microscope using ZEN pro software.

### 3D processing of the immunofluorescence images

In order to obtain the orthogonal cross-sectional analyses Z-stacks images were obtained and processed in Zen image processing software. The optimum slice size was 0.24 µm. The fluorescence intensities obtained were plotted in Sigma Plot 10.

### Immunoblotting

The immunobloting for Tau phosphoepitopes AT 8 and AT 100 was carried out for the same 6 treatment groups as immunofluorescence studies. 3 lakh cells were seeded in 60 mmm culture dishes and incubated for 24 hours in advanced DMEM supplemented with 10 % FBS and antibiotics. The treatment was given as described previously. After treatment cells were harvested and lysed in RIPA buffer. The lysate was centrifuged at 14000 rpm for 20 minutes and supernatant was separated and processed for protein estimation. 75 µg of protein was loaded onto 10 % SDS-PAGE and transferred onto PVDF membrane. The membranes were blocked in 5% BSA and incubated in primary antibodies AT 8 (1:1000) and AT 100 (1:1000) overnight at 4 °C. The blots were washed in PBST thrice and incubated with secondary HRP cross-linked antibodies for 1 hour. The unbound antibody was washed off by PBST and the blots were developed by enhanced chemiluminescence kit (ECL plus).

### Drosophila melanogaster Stock

*Drosophila Melanogaster* recombinant line-Elav GMR h Tau, that expresses the 2N4R isoform of human tau (MAPT) in the eye under the control of GMR driver, was obtained from Bloomington Drosophila Stock Center.

#### Fly Stock Maintenance

Flies were raised on standard Cornmeal Food [corn flour (75g), sugar (80g), yeast (24g), agar (10g), malt (60g), methyl paraben (5ml) and propionic acid (5ml) for 1Lt] and maintained in incubator (MODEL: DR-36VL, PERCIVAL) under a 12h light/12h dark cycle at 25 °C and 52% humidity.

### Toxicity assay and feeding

Four to five days old flies were fed on different concentrations of compounds (2 µM to 400 µM, depending on the compound). Mortality was recorded every 48 hr for 31 days. The concentrations at which no mortality was observed during this period were selected for further study to understand the protective efficacy of the compounds. Control flies were fed on water, DMSO (24µl) and ethanol (50,100 & 500 µl), depending on the solubility of the compounds.

### Survival Assay

Control and experimental flies were collected 24 h after hatching and maintained in incubator (MODEL: DR-36VL, PERCIVAL) on a standard cornmeal medium at 25 °C and 52% humidity in a 12:12 h light-dark cycle. Highest concentration of the compounds were used in the experiment as no toxicity or mortality was observed during the toxicity assay. Twenty five flies for each compound were maintained in a Drosophila vial. Flies were transferred to a fresh medium twice a week. Viability of flies for each compound was analysed daily. Survival of the flies along with their respective controls was recorded for 31 days. The data is presented in the form of segmented bar graph reflecting the percentage of live and dead flies observed at the end of the assay between experimental and control variants.

### Qualitative Assessment of Eye Phenotype

For light microscope imaging of adult eyes, 2 to 3-days old flies were decapitated and then imaged using a Nikon SMZ 100 Stereo Microscope. Images were captured with Nikon DS camera and the slices were stacked using NIS Elements D software.

### Statistical analysis

All the graphs were plotted in Sigma Plot 10. Unpaired t-test was performed to get the statistical significance. The error bars represent mean ±SD values with 95% confidence intervals.

## Declarations

### Ethics approval

Not applicable

### Consent to participate

Not applicable

### Availability of data and material

Not applicable

### Code availability

Not applicable

### Author contributions

SC designed the project. SKS and SC conducted the cell biology experiments, analyzed the results, and wrote the paper. SC conceived the idea for the project, supervised, resource provided and wrote the paper. AR and AM carried out the Drosophila related work.

